# PEGylation strategies for enhanced nanoparticle delivery to tumor associated immune cells

**DOI:** 10.1101/2025.07.23.666401

**Authors:** Devorah Cahn, Sanjay Pal, Nimit L. Patel, Timothy Gower, Senta M. Kapnick, Christopher M. Jewell, Gregg A. Duncan, Matthew T. Wolf

## Abstract

Barriers to nanoparticle drug delivery to the tumor microenvironment such as ECM deposition and clearance by the mononuclear phagocyte system have necessitated strategies for more effective tumor penetration. Adding polyethylene glycol (PEG) chains to the surface of nanoparticles (PEGylation) has been widely used to both enhance accumulation at the tumor site and increase blood circulation time. Recent work has also shown that immune cells (e.g. macrophages, dendritic cells, neutrophils) play an important role in the ability of NPs to effectively target and spread within a tumor. PEG chain characteristics such as size and branching affects how nanoparticles interact with tissues; however, it is unclear how PEGylation type affects NP uptake and cellular distribution in the tumor microenvironment. In this study, we evaluated the influence of both linear and branched PEGylation on nanoparticle biodistribution and uptake in tumor cells as well as tumor-infiltrating immune cells. As compared to conventional surface coatings with linear PEG, we show that modifying PEG structure to a branched conformation increases nanoparticle accumulation in the spleen of tumor-bearing mice, primarily due to significantly enhanced uptake by leukocytes. As compared to uncoated particles, we also found that nanoparticles densely coated with linear or branched PEG accumulated to a greater extent in tumors showing ≥8-fold increases in uptake by tumor-associated macrophages and dendritic cells. These studies provide insight into PEG architecture as a design parameter in nanomedicine that can facilitate the design of more effective cancer therapies.

**Translational Impact:** This work uncovers immune-mediated mechanisms and design strategies to enhance NP delivery to tumors via PEGylation which provide a foundation for clinical development of cancer therapeutics. The PEGylation strategies described could be readily integrated into clinically relevant nanoparticle delivery systems (e.g. lipid nanoparticles) with minimal effort making this a highly appealing approach to address the current limitations of cancer nanomedicine.

## Introduction

Nanoparticle (NP) drug carriers are often employed to effectively increase drug bioavailability and targeting to the tumor microenvironment (TME) in cancer (1–3). However, efficient delivery is often hindered by clearance through the mononuclear phagocyte system (MPS) in the spleen, liver, and blood (4–7). To circumvent this, NPs are often coated with polyethylene glycol (PEG) to shield their surfaces from serum protein adsorption (e.g. opsonins) and endow stealth function to NPs capable of evading the MPS-mediated clearance enabling longer circulation time (8). NPs must also possess the ability to cross tumor blood vessels. Passive transport through ‘leaky’ tumor vasculature via the enhanced permeability and retention (EPR) effect has been thought to facilitate NP transport into tumors but recent studies have questioned if this mechanism will translate across multiple cancer types (9–11). More recent work has identified endothelial and immune cell subtypes that participate in active NP transport into the TME (12). Once NPs arrive at the tumor site, they must also be able to efficiently travel through the ECM to reach and distribute within solid tumors to exert a therapeutic effect (13,14). Prior research by ourselves and others have demonstrated how PEGylated NP can more efficiently diffuse through the ECM enabling greater distribution within interstitial tissue regions (15–17). To successfully breach these biological barriers for effective NP delivery in cancer and other applications, there are several PEGylation design parameters worth considering, such as PEG density, chain length, and degree of branching. High-density PEG coatings have proven effective at enhancing NP circulation time as well as biodistribution in ECM-rich tissues such as the brain and solid tumors (18–20). More recently, we have found branched PEGylation may further improve diffusion and cellular uptake of NPs in a dense ECM microenvironment *in vitro* (16).

Immune cells play an essential role in the ability of NPs to effectively target and spread within the tumor microenvironment. Phagocytes in our innate immune system, such as macrophages, dendritic cells (DCs), and neutrophils, are the most abundant leukocytes in solid tumors where they influence tumor progression and the response to cancer therapy. Rather than directly targeting NP uptake to cancer cells, tumor phagocytes can be effective drug reservoirs, trapping and enriching drug-laden NPs within tumors (21,22). Neutrophils have been utilized for targeting NPs to the TME since they gravitate to inflammatory sites such as the TME and are the most abundant leukocyte found in the bloodstream (23). Studies have shown that neutrophils can effectively be used to penetrate the blood brain barrier and enhance nanodrug targeting in glioblastoma (24). Lin et al. showed that macrophages play an important role in tumor penetration where NPs penetrated more deeply into tumors when taken up by macrophages after extravasation (21). Furthermore, immune cells themselves are increasingly relevant targets for cancer immunotherapy. DCs are antigen-presenting cells that coordinate adaptive immune activation and are frequently targeted for immunotherapies (25,26). NPs are under investigation to deliver a variety of tumor antigens, immune stimulating adjuvants, and immune checkpoint blockade drugs to DCs (27). Phagocytes are therefore both targets for immunotherapy as well as active participants in particle retention and distribution within the TME.

As mentioned, PEGylated nanomedicine possesses stealth properties by design which is meant to limit their interactions with phagocytes. As such, PEGylated NP delivered systemically as vehicles for cancer therapeutics must strike a fine balance between evading clearance via immune recognition while remaining efficiently uptaken by tumor-associated immune cells for active trafficking into the TME. It is currently unclear to what extent PEGylation and resulting NP surface properties influence NP uptake by tumor-associated immune cells and immune cell-mediated NP transport into tumors. In addition to systemic infusion, direct intratumoral delivery of NP, while bypassing MPS mediated clearance, may also be limited by the dense ECM within the TME and this require strategies to enhance NP diffusion through this barrier to delivery. Therefore, in this study we examined whether dense coatings of linear and branched PEG can enhance NP accumulation within tumors and whether it affects immune cell infiltration and NP uptake in these cells. We compared both intravenous and intratumoral delivery of PEGylated NP biodistribution and uptake in the TME in both mouse and human xenograft tumor models. Moreover, both immune-competent and immune-deficient mice were evaluated to determine impact of these functions on delivery efficiency of PEGylated NPs. These studies will enable optimization of nanoparticle design strategies for tumor and immune cell targeting.

## Materials and Methods

### Materials

Fluorescent NIR carboxylate-modified polystyrene (PS-COOH) nanoparticles and live/dead fixable blue dye were obtained from Thermo Scientific. 5 kDa linear methoxy PEG-amine was purchased from Creative PEGWorks and 10 kDa 4-arm PEG amine was obtained from JenKem Technologies. Athymic nude and Balb/C? mice were obtained from the Jackson Laboratory.

### Nanoparticle preparation

5 kDa linear PEG or 10 kDa 4-arm branched PEG were coated on carboxylate-modified 100 nm red or NIR fluorescent polystyrene (PS) nanoparticles (PS-COOH) via a carboxyl-amine linkage as described in our previous work (15,16). Nanoparticle size and zeta potential were measured in 10 mM NaCl at pH 7 using a Nanobrook Omni (Brookhaven Instruments). Nanoparticle concentration was determined by fluorescence intensity by serially diluting stock NPs of known concentration to create a standard curve to calculate NP concentration in each group.

### Cell lines

The RAW 264.7 murine macrophage cell line was obtained from ATCC (# TIB-71, RRID: CVCL_0493) and confirmed negative for mycoplasma using MycoStrip (InvivoGen) at least twice prior to use. RAW 264.7 cells originated from a male mouse tumor and has not been previously reported as misidentified or contaminated. Additional authentication was not performed. A549 human lung cancer cell line was obtained from Division of Cancer Treatment and Diagnosis (DCTD) Tumor repository NCI (# CRM-CCL-185, RRID:CVCL_0023) which were authenticated by the DCTD. A549 cells were originally isolated from the lung of a 58-year-old, White male with carcinoma and has not been previously reported as misidentified or contaminated. 4T1 murine triple negative breast cancer cells were obtained from ATCC (# CRL-2539, RRID:CVCL_0125). 4T1 cells originated from female BALB/c mouse mammary tissue and has not been previously reported as misidentified or contaminated. Both A549 and 4T1 cells underwent Molecular Testing of Biological Materials (MTBM) by the NCI Frederick LASP Animal Diagnostic Laboratory prior to *in vivo* studies to confirm they were negative for mycoplasma and a panel of murine viruses: mouse hepatitis virus, polyoma virus, sendai virus, pneumonia virus of mice, reovirus 3, minute virus of mice, Theiler’s murine encephalomyelitis virus, lymphocytic choriomeningitis virus, Ectromelia virus, Lactic dehydrogenase-elevating virus, mouse parvovirus, mouse norovirus, mouse rotavirus, mouse adenovirus, and mouse cytomegalovirus.

### Bone marrow derived dendritic cells (BMDC) isolation, cell culture, and nanoparticle uptake

Femurs were harvested from male 8-12-week-old C57BL/6J mice under sterile conditions and the bone marrow flushed with cold Hanks’ Balanced Salt Solution (HBSS) and filtered through a 70um strainer. Red blood cells were lysed using RBC lysis buffer and cell suspensions were washed before counting. 1×10^6^ cells/mL were cultured in 10mL complete RPMI 1640 (RPMI-1640 supplemented with 10% heat-inactivated fetal bovine serum (FBS), 2mM L-glutamine, 55mM b-mercaptoethanol, 100U Pen/Strep; cRPMI) per 100mm petri dish in the presence of 20 ng/mL of granulocyte macrophage-colony stimulating factor (GM-CSF). On day three, fresh cRPMI supplemented with GM-CSF was added to cultures. On day six, a 1:1 v/v exchange of supplemented cRPMI was performed. Gentle pipetting in cold PBS was performed to remove cells between 7-9 days after the start of cultures for downstream assays.

BMDCs were seeded in a 96-well plate at a density of 100,000 cells per well. Cells were incubated at 37°C and 5% CO_2_ for 2 hours until cells adhered to the surface of the wells. Media was removed from on top of the cells in each well and replaced with 100 μL of media containing red fluorescent NPs at a concentration of 10^11^ NPs per mL. The cells were incubated with the NPs for 2 hours at 37°C and 5% CO_2_. After incubation, the NP containing media was removed from the cells and the cells were washed 3 times with cold PBS. The cells were then incubated with cold 2 mM EDTA in PBS for 5 minutes at 4°C to detach them from the wells and 4 wells were combined for each replicate for each group. Cells were then collected via centrifugation at 300 xg for 5 minutes and washed with cold PBS. To stain for dead cells, Zombie NIR dye was diluted 1:1000 and 300 μL was added to each and incubated for 25 minutes at 4°C. Cells were then washed twice with cold PBS by centrifuging at 300xg for 5 minutes and then fixed with 4% paraformaldehyde (PFA) for 15 minutes at 4°C. Cells were again washed twice with cold PBS via centrifugation at 300xg for 5 minutes. The cells were then resuspended in PBS and NP uptake was analyzed using a BD FACSCelesta flow cytometer.

### RAW 264.7 macrophage cell culture and nanoparticle uptake

RAW 264.7 macrophages obtained from ATCC were seeded on plastic in DMEM medium supplemented with 10% FBS and 1% penicillin-streptomycin and incubated at 37°C and 5% CO_2_. Cells were passaged upon reaching 70-80% confluency at which time they were dissociated from the plate surface using a cell scraper. Cells were seeded in a 24-well plate at a density of 500,000 cells per well and incubated at 37°C and 5% CO_2_ for 24 hours until they adhered to the plate surface. The media on top of the cells was then replaced with 500 μL of media containing red fluorescent NPs at a concentration of 10^11^ NPs per mL. The cells were then incubated with the NPs for 2 hours at 37°C and 5% CO_2_. After incubation, the NP containing media was removed from the cells and the cells were washed 3 times with cold PBS. Cell scrapers were used to detach the cells and cells were collected via centrifugation at 300 xg for 5 minutes. Dead cells were stained with 300 μL of Zombie NIR dye diluted 1:1000 for each replicate and incubated for 25 minutes at 4°C. Cells were then washed twice with cold PBS by centrifuging at 300xg for 5 minutes and then fixed with 4% paraformaldehyde (PFA) for 15 minutes at 4°C. Cells were washed twice with cold PBS via centrifugation at 300xg for 5 minutes, resuspended in PBS, and NP uptake analyzed using a BD FACSCelesta flow cytometer.

### Animal care

5-week old female balb/c and athymic nude mice were obtained from The Jackson Laboratory housed at the NCI Frederick laboratory Animal Sciences Program under 12-hr light/dark cycles in specific pathogen-free conditions. Animals acclimated for 2-3 weeks prior to use at 8-weeks old. The Animal Care and Use Committee at NCI Frederick (Protocol No. 20-063) provided ethical approval for animal experiments. NCI-Frederick is accredited by AAALAC International and follows the Public Health Service Policy for the Care and Use of Laboratory Animals. Animal care was provided in accordance with the procedures outlined in the ‘Guide for Care and Use of Laboratory Animals (National Research Council; 1996; National Academy Press; Washington, D.C.). Mice were euthanized via asphyxiation with carbon dioxide and cervical dislocation as the secondary method.

### NP administration and imaging in 4T1 and A549 tumors

Mouse mammary 4T1 cells were cultured with RPMI medium supplemented with 10% FBS and 1% penicillin-streptomycin at 37°C and 5% CO_2_. Human lung epithelial A549 cells were seeded on plastic in Ham’s F-12K (Kaighn’s) medium supplemented with 10% FBS and 1% penicillin-streptomycin and incubated at 37°C and 5% CO_2_. Both cell lines were passaged upon reaching 70-80% confluency at which time they were dissociated from the plate surface using 0.05% trypsin EDTA for 5 minutes at 37°C. Cells were washed 3 times in 1X PBS and resuspended in 1X PBS to a final concentration of 5×10^6^ cells/mL and 1×10^7^ cells/mL for 4T1 and A549 cells respectively. 100 μL of the cell suspension was subcutaneously injected into the right flank of 8-week-old athymic nude mice. 100 μL of the 4T1 cell suspension was also subcutaneously injected into the right flank of 5-week-old balb/c mice. Tumors were grown to a size >5 mm in greatest diameter (9 days or 4 weeks for 4T1 or A549 tumors, respectively), after which mice were intravenously injected via the tail vein with 100 ul of PEG-L5, PEG-B10, or non-PEGylated NPs at a concentration of 10^13^ NPs/mL (10^12^ NPs/mouse) or with 100 uL of PBS as vehicle control. An IVIS Spectrum (Perkin-Elmer) was used for live animal fluorescence imaging immediately after injections as well as at 6, 24, 48, and 120 hours later. A randomized cohort from each treatment group was euthanized 24 hours post injection, and the tumor, liver, and spleen from each mouse was harvested and imaged *ex vivo* using IVIS. The total radiance efficiency was calculated for each organ by normalizing the fluorescence of the region of interest (ROI) to the ROI area.

### Cell isolation and immunostaining

Cells were isolated from harvested tissue by dicing the tissue on ice and digesting the tissue using 0.25 mg/mL Liberase TL and 0.2 mg/mL DNAse I in 5 mL RPMI for 20 minutes at 37°C on a shaker. The mixture was then passed through a 70-um cell strainer using the back of a syringe plunger and cold 1X PBS. The collected cells were centrifuged at 300 xg for 5 minutes at 4°C and the pellet was washed with cold PBS. 5 mL of 1x RBC lysis buffer was added to the cells for 3 minutes on ice and then washed twice with cold PBS. Viability staining to exclude dead cells was performed with live/dead blue fixable dye diluted 1:1000 in PBS and incubated with cells for 25 minutes at 4°C followed by washing twice in cold PBS. Cells were then stained with anti-CD45 (BUV395), anti-CD11b (AF700), anti-Ly6G (eFluor450), anti-CD11c (BUV805), anti-F4/80 (BV785), and anti-CD86 (AF647) with Fc block and monocyte block in 100 uL of FACS buffer (0.5% BSA in 1X PBS) for 40 minutes at 4°C and washed twice with cold FACS buffer by centrifuging at 300 xg for 5 minutes at 4°C. The cells were resuspended in FACS buffer and analyzed using a 5-laser Cytek Aurora Spectral Flow Cytometer and SpectraFlo version 3.3.0 software. Flow cytometry data was further analyzed and displayed using FlowJo software (version 10.10.0, BD Biosciences).

### Statistical Analysis

GraphPad Prism 9 (GraphPad Software) was used for graphing and statistical analysis of the data. A two-way analysis of variance (ANOVA) with a Tukey post-hoc test for analysis of differences between groups. Bar graphs show mean and standard deviation. Statistical significance was assessed at *p* < 0.05.

## Results

### PEGylation modulates NP uptake in BMDCs and macrophages in vitro

PEGylation is a widely used strategy to delay NP removal from circulation by immune cells. Previous studies have shown that branched PEG conformations can increase blood circulation time compared to their linear counterparts potentially due to an increased shielding effect (28,29). However, it is unclear whether and to what extent PEG branching affects NP uptake in different types of immune cells *in vitro*. To determine whether PEG branching can decrease NP uptake in different phagocyte populations, we incubated macrophages and dendritic cells with NPs coated with either 5 kDa linear PEG (PEG-L5) or with 10 kDa 4-arm branched PEG (PEG-B10) compared to uncoated non-PEGylated NPs (No PEG). The linear PEG conformation was chosen to match the fully extended length of the branched PEG from the surface of the NPs (15). After PEG conjugation, we found both PEG-L5 and PEG-B10 NP possessed near neutral surface charges (zeta potential of ∼0 mV) and similar diameters in the 125-150 nm range (**Figure S1**). Both types of PEGylated NPs reduced the percentage of BMDCs that internalized NPs relative to non-PEGylated NPs (64.1 ± 2.1%), with a stronger reduction observed for PEG-L5 NPs (2.0 ± 1.8%) compared to the PEG-B10 NPs (15.2 ± 12.1%) (**Figure 1A**). We estimated the relative amount of NP uptake per cell from the geometric fluorescence intensity (gMFI) and found it was also significantly lower for PEGylated NPs compared to non-PEGylated NPs (**Figure 1B**). NP uptake per cell was 8.6-fold lower for PEG-B10 and 14.1-fold lower for PEG-L5 relative to non-PEGylated NPs. Though the difference between PEG-B10 and PEG-L5 was statistically significant, both largely abrogated NP uptake under *in vitro* conditions. A similar effect was observed for NP uptake in RAW 264.7 macrophages (**Figure 1C-D**) where PEG-L5 NPs prevented uptake, both as a percentage of cells and uptake per cell. However, PEG-B10 did not strongly reduce the percentage of cells internalizing particles showing phagocyte specific responses to branched PEGylation (**Figure 1C-D).**

**Figure 1.**
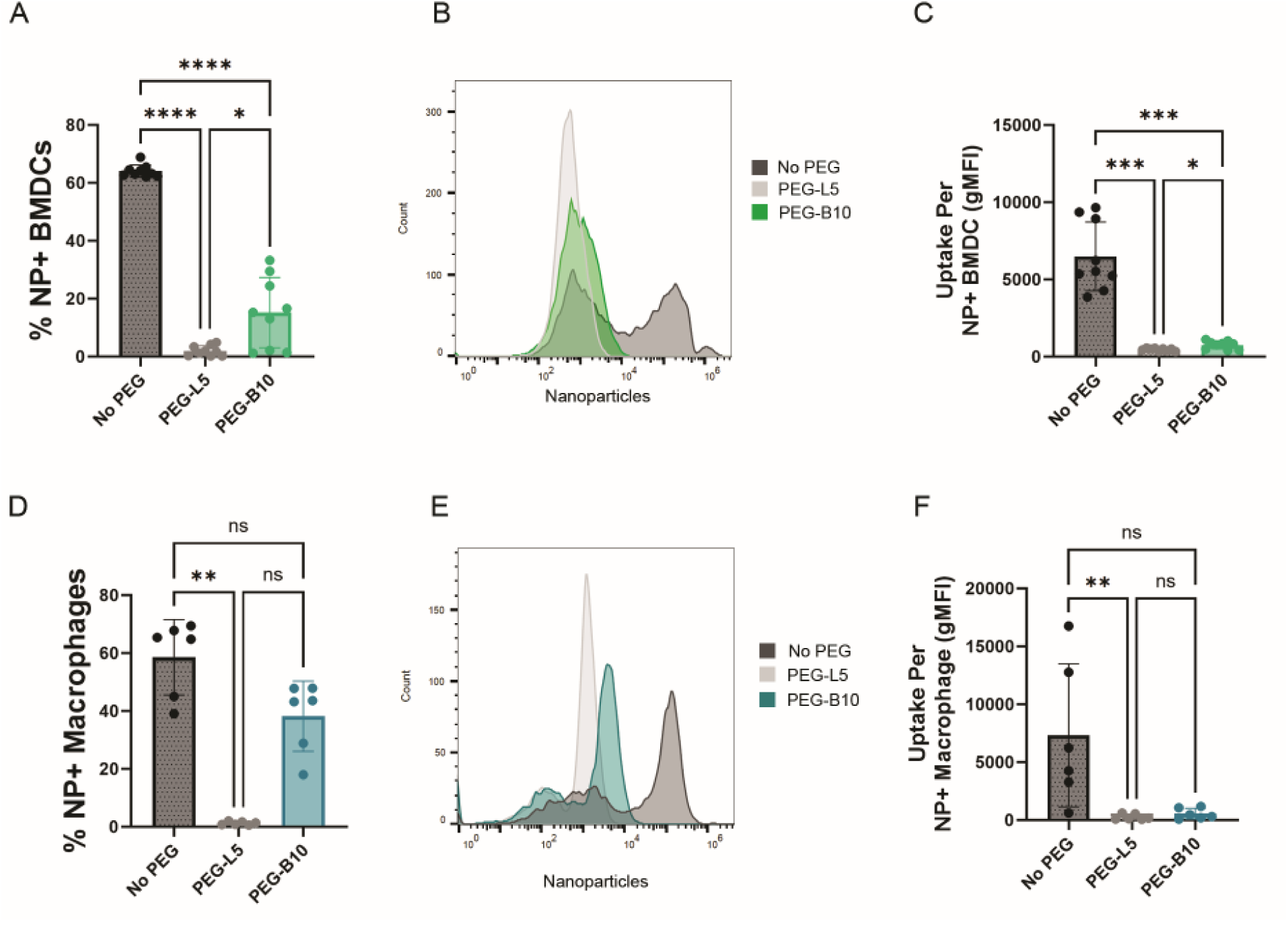
Nanoparticle uptake in phagocytes *in vitro*. (A) Percent of BMDCs internalizing NPs. (B) Representative histograms of the fluorescence intensity of NPs in BMDCs. (C) Geometric mean fluorescence intensity (gMFI) of NPs in BMDCs. (D) Percent of RAW 264.7 macrophages internalizing NPs. (E) Representative histograms of the fluorescence intensity of NPs in RAW 264.7 macrophages. (F) gMFI of NPs in RAW 264.7 macrophages. *p<0.05, **p<0.01, ***p<0.001, and ****p<0.0001 by two-way ANOVA with Tukey correction.

### PEGylation increases NP spreading after intratumoral injection

Intratumoral (IT) injection is a strategy to locally deliver high concentrations of NP therapeutic payloads within tumor tissue without systemic exposure to the MPS or the need to pass endothelial barriers. However, intratumoral injection is often difficult to translate in clinical settings where it requires guidance using medical imaging. Nonetheless, intratumoral administration enables direct assessment of how surface chemistry influences the spread of NPs within the tumor interstitial space.

To determine whether PEGylation and PEG architecture affect NP distribution throughout the tumor, we cryosectioned 4T1 tumors 24 hours post IT injection and took images of the sections using a fluorescent microscope to visualize NP spread. The tumors injected with non-PEGylated NPs had little visible NP fluorescence throughout the tumor with only a small portion of the tumor edge exhibiting significant NP fluorescence, likely from the injection site (**Figure 2**). PEG-L5 and PEG-10 NPs were similarly well distributed in the 4T1 model, though we found more consistent tumor coverage in the A549 xenograft with PEG-B10 NP. These findings suggest that PEGylation influences spreading through tumor tissue and not just increasing half-life in circulation by evading MPS clearance. Similarly, we cryosectioned A549 tumors 24 hours post IT injection and took images of the sections using a fluorescent microscope to visualize NP spread. Results were comparable to what was visualized for NP spread in 4T1 tumors. The tumors injected with non-PEGylated NPs had little visible NP fluorescence throughout the tumor with only a small portion of the tumor edge exhibiting significant NP fluorescence, likely from the injection site (**Figure 2**). Tumors treated with both PEG-L5 and PEG-B10 NPs had substantial visible NP fluorescence spread throughout the tumor (**Figure 2**).

**Figure 2.**
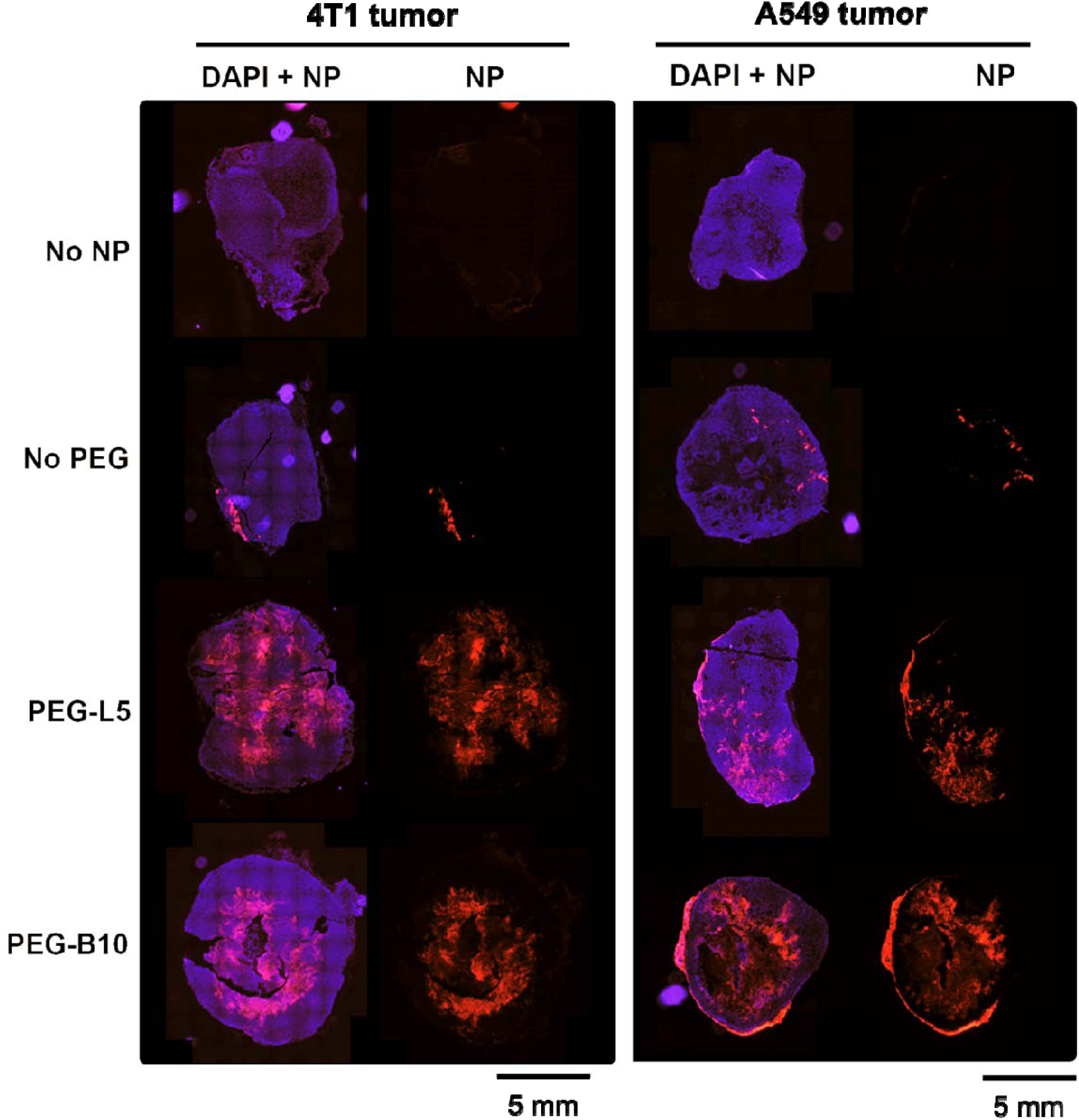
4T1 and A549 tumor section fluorescent micrographs after intratumoral NP injection. Representative images of serial tumor cryosections that were either untreated or intratumorally injected with non-PEGylated, PEG-L5, or PEG-B10 NPs 24 hours before harvest in nude mice (N=3/group).

### Branched versus linear PEGylation differentially alters NP biodistribution after intravenous delivery in 4T1 tumor bearing athymic nude mice

Since we found both linear and branched PEGylation improved NP distribution within the TME with intratumor delivery, we further assessed the biodistribution of PEG-L5 and PEG-B10 coated NP following systemic delivery in tumor bearing mice. Previous work has shown that PEGylation can increase NP circulation time and increase NP biodistribution in tumor tissue *in vivo*, increasing overall NP uptake in tumor target cells (8,28). Recent studies have also shown that PEG branching can reduce the accelerated blood clearance (ABC) phenomenon by preventing binding to anti-PEG antibodies that develop with persistent exposure to PEGylated carriers (30). Our previous research also showed that PEG branching increases NP mobility in the extracellular matrix (ECM) suggesting it may improve penetration through fibrous tumors (15,16). Therefore, we examined how PEG branching affected NP accumulation and retention at the tumor site as well as in tissues important for clearance. Hematopoiesis and phagocyte function are systemically modulated during tumor progression; therefore, we quantified system-wide distribution in tumor bearing animals.

Athymic nude mice readily formed subcutaneous 4T1 breast tumors, reaching ∼1 cm in diameter in 9 days (**Figure S2, S3**). Mice were injected with either PEG-L5, PEG-B10, or non-PEGylated NPs or with saline as a control at a total dose of 10^12^ NPs. Tumor growth was not affected by NP administration in any group (**Figure S3**). Live animal imaging showed that the type of PEGylation altered NP distribution within the tumor as well as in the spleen and liver, which are major sites of MPS clearance (**Figure 3A, B**). All PEGylated NPs spread broadly throughout the body while non-PEGylated NPs generally accumulated in the center, which is consistent with anatomic location of the liver and spleen (**Figure 3A**). NP accumulation within 4T1 tumors most rapidly accumulated within 24 hours of injection, reaching steady state by 48 hours that persisted to the 5-day study endpoint (**Figure 3B**). Tumor uptake was similarly increased by ∼35-fold for PEG-B10 and ∼14-fold for PEG-L5 relative to non-PEGylated NPs at 24 hours (**Figure 3C,D**). Non-PEGylated NPs accumulation was greatest in the liver followed by spleen, and PEG-L5 reduced liver accumulation by approximately half. (**Figure 3C-D**). PEG-B10 had the greatest enrichment in *ex vivo* tumors, though surprisingly, did not reduce liver signal and increased spleen uptake.

**Figure 3.**
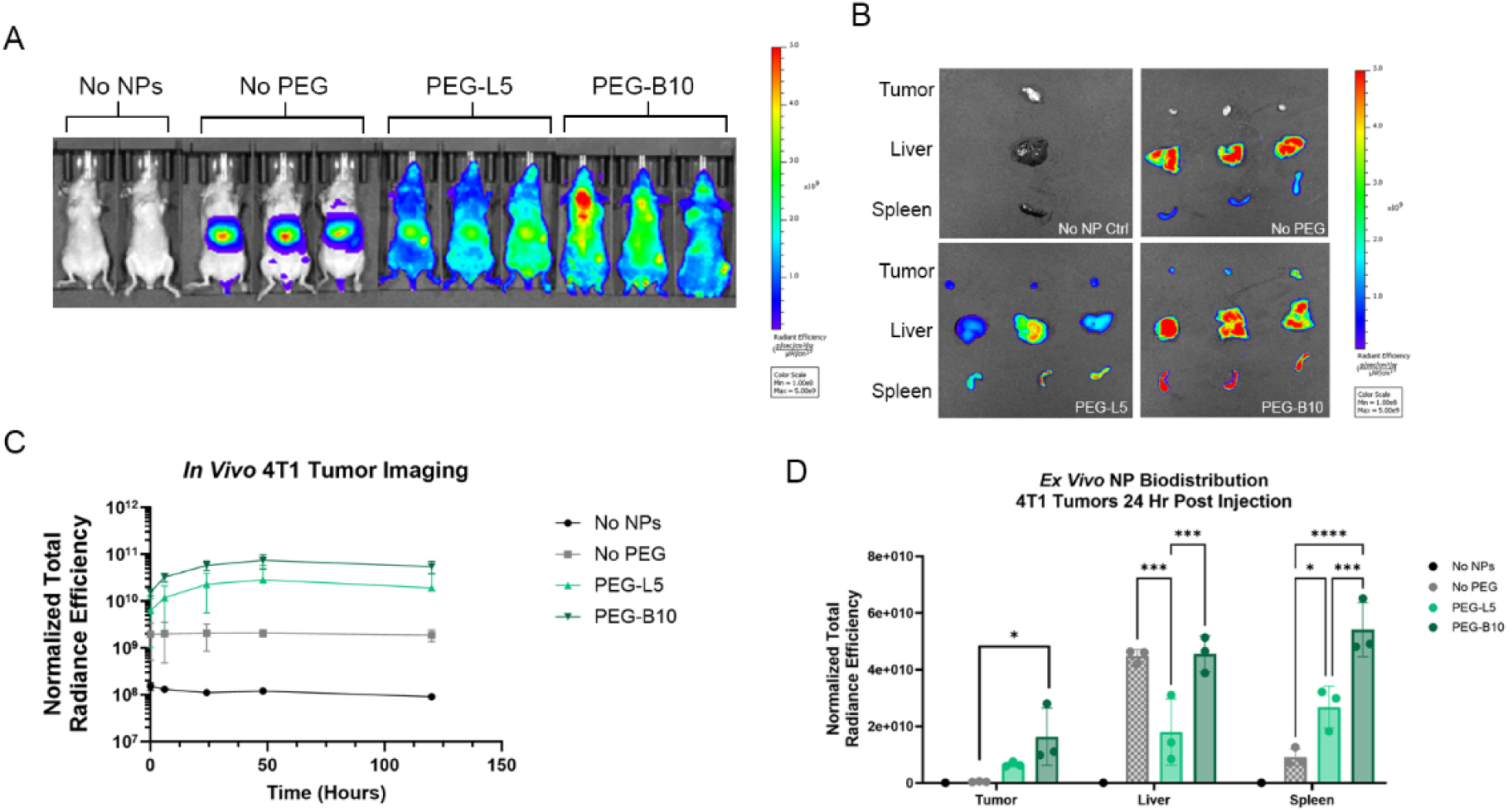
In vivo and ex vivo NP biodistribution in the 4T1 tumor model in nude mice. **(A)** Live animal imaging of NP biodistribution. **(B)** Ex vivo imaging of the tumor, liver, and spleen of mice in different treatment groups. **(C)** Analyzed total radiance efficiency of the tumors in mice for different treatment groups. **(D)** Analyzed total radiance efficiency of ex vivo images of the tumor, liver, and spleen from different treatment groups. **p<0.01 and ***p<0.001 by two-way ANOVA with Tukey correction, N=3 mice for No PEG, PEG-L5, and PEG-B10, N=1 mouse for No NPs.

### Branch PEGylation enhances NP uptake in leukocytes in 4T1 tumors after intravenous delivery in nude mice

We then performed flow cytometry analysis to define which cell types were responsible for the observed NP accumulation within 4T1 tumors with an emphasis on phagocytes given their well-established role in nanoparticle delivery to solid tumors. For example, prior work has shown that NP distribution within the tumor microenvironment is directly tied to their uptake by macrophages, and that NP laden macrophages penetrated deeper into tumors (21). Others have utilized the tendency of neutrophils to accumulate at the tumor site to enhance NP tumor distribution while dendritic cells are commonly targeted in immunotherapy applications (23,24,31). We determined which phagocyte and non-immune cell populations internalized branch PEGylated NPs within the tumor microenvironment as well as in the liver and spleen (**Figure 4**). It should be noted that athymic nude mice lack mature T cells and as such, we did not include this population in our analysis. The immune status in this model would mirror the situation for patients undergoing cytotoxic chemotherapy and T cell ablation prior to CAR T-cell therapy (32). Nevertheless, we do expect the lack of functional T cells influenced NP biodistribution in this model and considering this, we performed subsequent studies in mice with a fully competent immune system included in this study.

**Figure 4.**
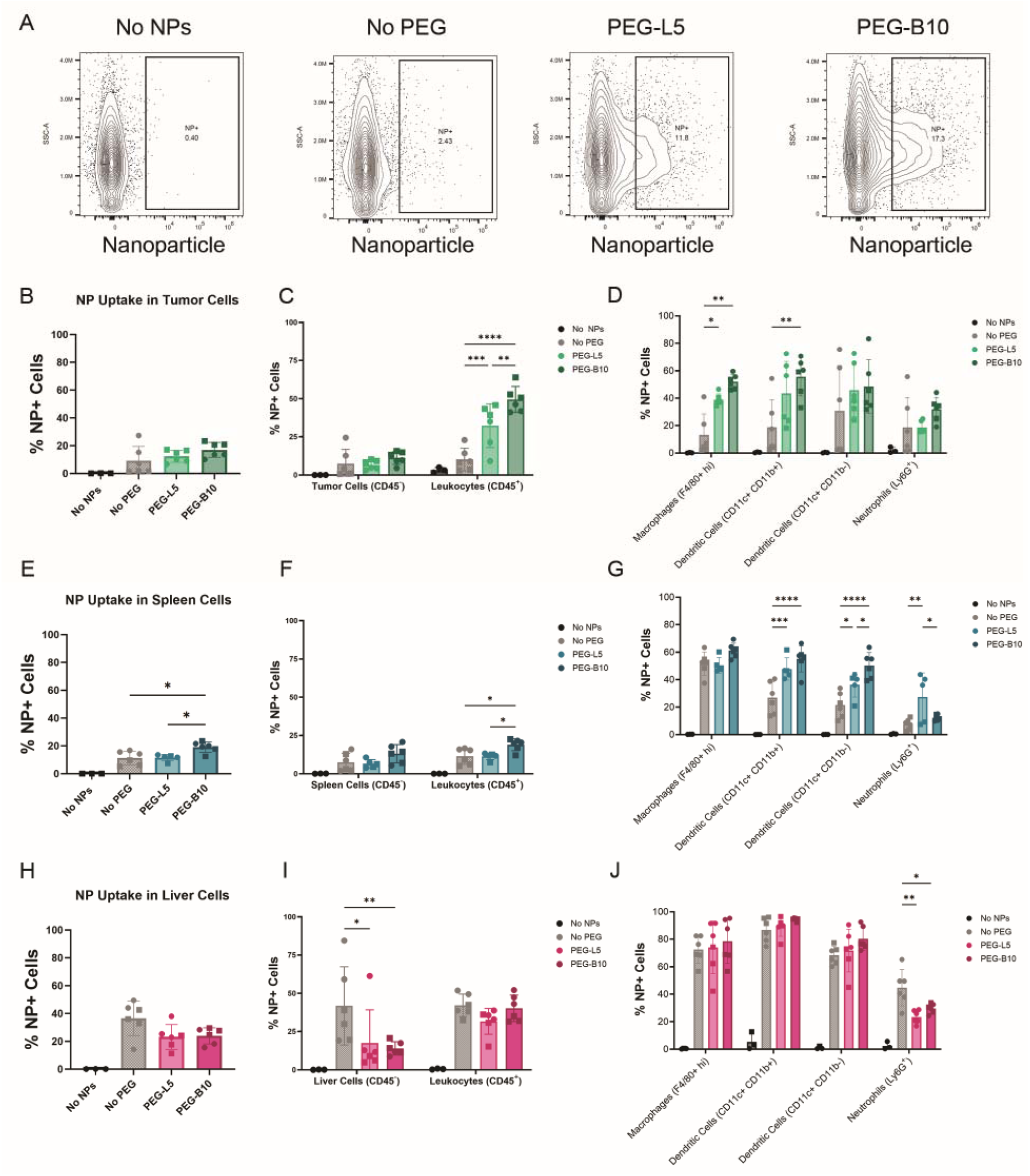
NP immune cell uptake in 4T1 tumor bearing nude mice. **(A)** Representative flow cytometr contour plots of NP gating strategy for live cells isolated from 4T1 tumors. Percent of **(B)** live cells, **(C)** leukocytes or tumor cells, **(D)** neutrophils, dendritic cells, or macrophages containing NPs obtained from tumor tissue. Percent of **(E)** live cells, **(F)** leukocytes or CD45-spleen cells, **(G)** neutrophils, dendriti cells, or macrophages containing NPs obtained from spleen tissue. Percent of **(H)** live cells, **(I)** leukocytes or liver cells, **(J)** neutrophils, dendritic cells, or macrophages containing NPs obtained from liver tissue. *p<0.05, **p<0.01, and ***p<0.001 by two-way ANOVA with Tukey correction. N=6 mice for No PEG, PEG-L5, and PEG-B10, N=4 mice for No NPs

The overall percentage of 4T1 tumor cells in nude mice internalizing NPs did not substantially change with PEGylation, but it greatly affected the cellular composition internalizing cells. PEG-B10 NPs trended towards the highest tumor uptake overall (17.5% of live cells) 24 hours after injection compared to PEG-L5 NPs (12.4%) and non-PEGylated (9.1%) (**Figure 4A,B and Figure S5**). Of these live tumor cells, cancer and stromal cell uptake was similar across groups (1-5%), but there was a striking increase in CD45+ leukocyte uptake (**Figure 4C and Figure S5)** with PEGylation that was greatest in PEG-B10 (49.5%) and moderately increased in PEG-L5 (32.3%) compared to non-PEGylated (10.1%). Further leukocyte phenotyping determined that phagocytes were highly sensitive to NP PEGylation type and contributed to this increased uptake (**Figure 4D and Figure S5**). Macrophages (F4/80-high, CD11b+) showed the greatest increase in NP uptake with PEGylation, from 13% in Non-PEGylated to 51.9% with PEG-B10, and slightly lower effect in PEG-L5 (38.9%). NP uptake in DCs was also affected by PEGylation, especially monocyte derived DCs (CD11b+ CD11c+) which had the greatest NP uptake in PEG-B10 (55.6%) compared to non-PEGylated (18.7%). Classical DC (CD11b-CD11c+) uptake was consistently high in both PEG-L5 and PEG-B10 (45.7% and 48.3%, respectively), but was variable for Non-PEGylated NPs. In addition, neutrophils showed substantial uptake, which trended greatest with PEG-B10 (∼18% vs 31% for PEG-B-10). The majority of NP+ macrophages and DCs were CD86+. Neutrophils were not, and the F4/80+ macrophages co-expressed CD11c as part of this tumor associated macrophage phenotype (**Figure S4 and Figure S6A,B**). Additionally, other less characterized cell types demonstrated NP uptake, including both CD11b+ and CD11b-subsets that did not co-express the phagocyte markers in this panel (**Figure S4 and Figure S6A,B**). We also identified a Ly6G+ macrophage population that had similarly high uptake as conventional tumor macrophages.

We further performed this flow cytometry analysis on liver and spleen tissue which showed altered distribution profiles with PEGylation (**Figure S4 and S6C-F**). Within the spleen, PEG-B10 showed the overall greatest uptake in total live cells and in CD45+ leukocytes (**Figure 4E**). Of splenic phagocytes, macrophages had the highest percentage of NP positive cells (greater than 50%+ in all groups) followed by CD11b+ and CD11b-dendritic cells, while NP uptake within neutrophils was relatively low overall except in the PEG-L5 group which exceeded 20% (**Figure 4G**). PEGylation strategy had the greatest influence on NP uptake in DCs (both CD11b+/-), with PEG-B10 NPs consistently showing higher internalization over linear or non-PEGylated NPs. Consistent with the literature, liver tissue had the greatest overall uptake of the tissues tested (**Figure 4H**). Non-PEGylated NPs had the highest uptake (**Figure 4I**) which agreed with live animal imaging results (**Figure 3**), and was enriched in CD45-cells. NP uptake in macrophage and dendritic populations was unaffected by PEGylation. (**Figure 4J**) Interestingly, non-PEGylated NPs displayed enhanced accumulation in neutrophils, whereas PEGylated NPs were decreased. The 4T1 model is known to drive aberrant hematopoiesis and neutrophil maturation with greater frequencies in tissues such as the liver where they support metastasis (33).

### Branched and linear PEGylation alters NP biodistribution after intravenous delivery in A549 human lung tumor xenografts

Properties of the tumor microenvironment such as ECM content and tissue stiffness as well as immune cell infiltration are significantly impacted by cancer type (34). Therefore, we investigated how PEG branching would affect NP accumulation and immune cell infiltration within tumors in a human lung adenocarcinoma to determine if PEGylation effects were generalizable to a human cancer cell line. We subcutaneously injected athymic nude mice with A549 cells for injection with NPs once tumors reached a minimum diameter of 5 mm, which occurred within 28 days (**Figure S7, S8**). The same NP groups were compared (PEG-L5, PEG-B10, or non-PEGylated NPs or with saline as a control), and there were no notable differences between tumor volumes over the course of the study (**Figure S8A-C**). Twenty-four hours after NP injection, we harvested the tumor, spleen, and liver of each mouse and imaged them ex vivo (**Figure 5A**). Similar to the mouse 4T1 breast tumor model, non-PEGylated NPs had particularly high fluorescence intensity in the liver (**Figure 5A**), while both PEGylated NP formats were most abundant in the spleen followed by liver. NP signal was lowest in the tumor for all groups, though PEG-L5 NPs had slightly greater tumor accumulation (**Figure 5B**). This supports that the overall effect of PEGylation may be tumor type dependent.

**Figure 5.**
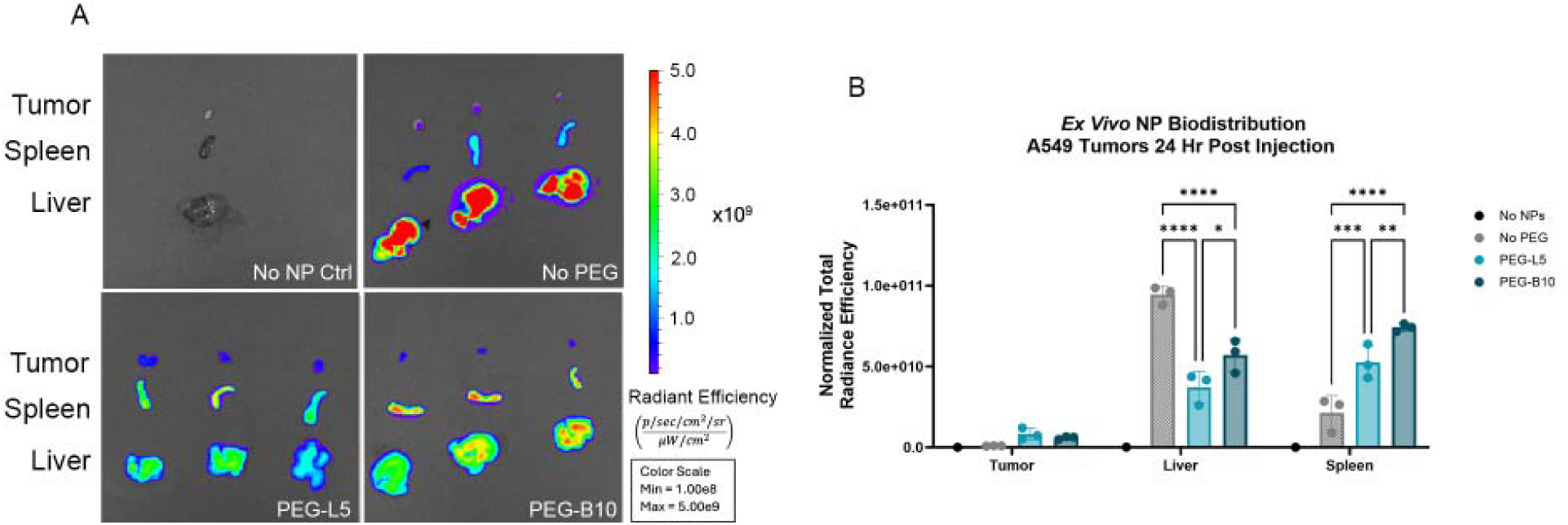
Ex vivo NP biodistribution in A549 tumor model. **(A)** Ex vivo imaging of NP biodistribution. **(B)** Analyzed total radiance efficiency of the tumors, spleen, and liver in mice for different treatment groups ex vivo. N=3 mice for No PEG, PEG-L5, and PEG-B10, N=1 mouse for No NPs. Mean and standard deviation is displayed with ANOVA and post-hoc Tukey test to determine significance.

### PEGylation alters cellular uptake distribution in A549 tumors after intravenous delivery in nude mice

Total NP uptake was increased with PEGylation within A549 human xenograft tumors, from approximately 1% in non-PEGylated to over 12% in PEG-L5 and PEG-B10. Unlike 4T1 tumors, there was negligible non-PEGylated accumulation in any A549 tumor cells, consistent with extremely low signal with IVIS imaging. Although total A549 tumor fluorescence was low in all NP groups via live animal imaging, we found PEGylation strategy altered immune cell biodistribution within the tumor microenvironment (**Figure S9A,B**). PEG-L5 NPs exhibited the highest uptake within CD45+ leukocytes followed by PEG-B10 NPs and very low overall uptake for non-PEGylated NPs (**Figure 6A-C**). Both PEGylation types enhanced macrophage and DC internalization compared to non-PEGylated. Macrophages and DCs were the most affected, with the greatest internalization with PEG-L5 (27%) followed by PEG-B10 (18%) in macrophages. Monocyte derived DC uptake was greater with PEG-L5 than PEG-B10 (**Figure 6D**) but was highest in classical DCs with PEG-B10.

**Figure 6.**
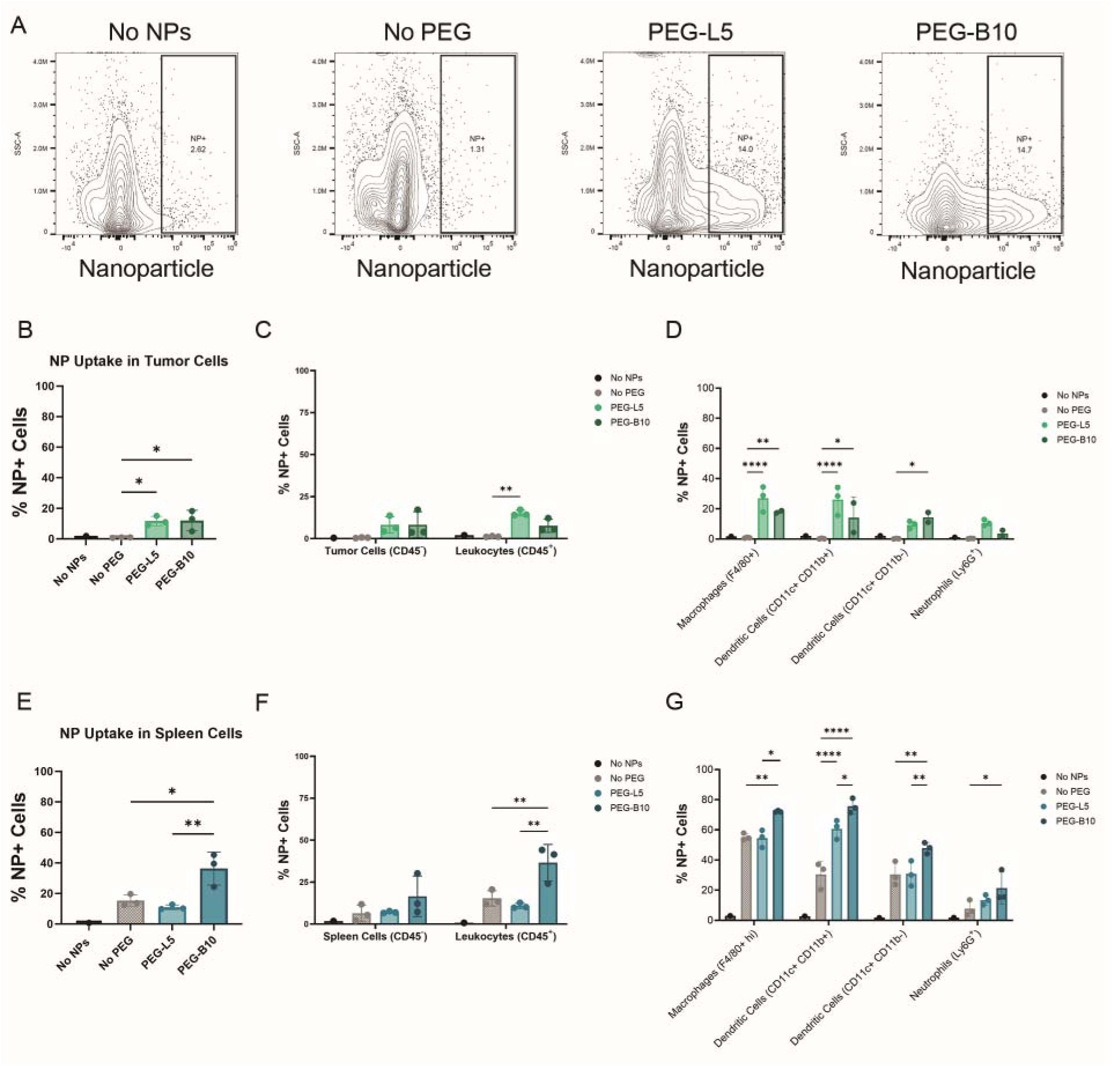
NP immune cell uptake in A549 tumor bearing nude mice. **(A)** Representative flow cytometry contour plots of NP gating strategy for live cells isolated from A549 tumors. Percent of **(B)** live cells, **(C)** leukocytes or CD45-tumor cells, **(D)** neutrophils, dendritic cells, or macrophages containing NPs obtained from tumor tissue. Percent of **(E)** live cells, **(F)** leukocytes or CD45-spleen cells, **(G)** neutrophils, dendritic cells, or macrophages containing NPs obtained from spleen tissue. *p<0.05, **p<0.01, and ***p<0.001 by two-way ANOVA with Tukey correction. N=3 mice for No PEG, PEG-L5, and PEG-B10, N=1 mouse for No NPs.

We analyzed spleens from A549 tumors and found PEG-B10 enhanced total NP accumulation, consistent with live animal imaging, and the majority of which were CD45+ leukocytes (**Figure 6F and Figure S9**). Indeed, all macrophage, DC, and neutrophil populations within spleens from A549 tumor bearing mice showed enhanced uptake compared to non-PEGylated NPs (**Figure 6G**). PEG-L5 only affected CD11b+ DCs. Like the findings within the 4T1 tumors, macrophages and dendritic cells within the spleen exhibited the highest NP uptake. Though these tumors were in the same strain of mouse, the NP uptake profile in the spleen differed by tumor type, which suggests systemic tumor related changes to NP biodistribution.

### Branched and linear PEGylation enhance NP uptake in immune competent 4T1 tumors after intravenous delivery

As previously noted, athymic nude mice lack mature T cells which help shape the TME, and dynamically regulate phagocyte function in tumors and other tissues. Furthermore, many patients receiving nanotherapeutics will not have had T cell ablation. Therefore, we isolated cells from 4T1 tumors implanted in fully immune-competent wild-type balb/c mice. There was similar overall uptake in tumors of mice treated both the PEG-L5 and PEG-B10 NPs (26.6% and 24%, respectively) which was up ∼5 fold greater than non-PEGylated NPs that had relatively little uptake (5%) (**Figure 7A and Figure S10A,B**). Unlike nude mice, PEGylation increased CD45-cell (cancer and stromal cells) uptake, from 2.3% in non-PEGylated to 13.4% and 14.5% in PEG-L5 and PEG-B10, respectively. Similar to 4T1 tumors in nude mice however, PEGylation caused a striking increase in CD45+ leukocyte uptake to 40-50% of cells. Macrophages were the greatest contributor to this increase, which was below 20% in non-PEGylated increasing to over 70% of cells in both PEG-L5 and PEG-B10 groups. Significant uptake of PEGylated NPs were also observed in monocyte derived DCs (up to 54%, ∼5-fold) and in classical DCs (up to 37%, ∼8-fold) compared to non-PEGylated. Neutrophils were not significantly affected. Generally, PEG-L5 and PEG-B10 showed nearly identical increases in NP uptake within 4T1 tumors implanted in immune competent balb/c mice.

**Figure 7.**
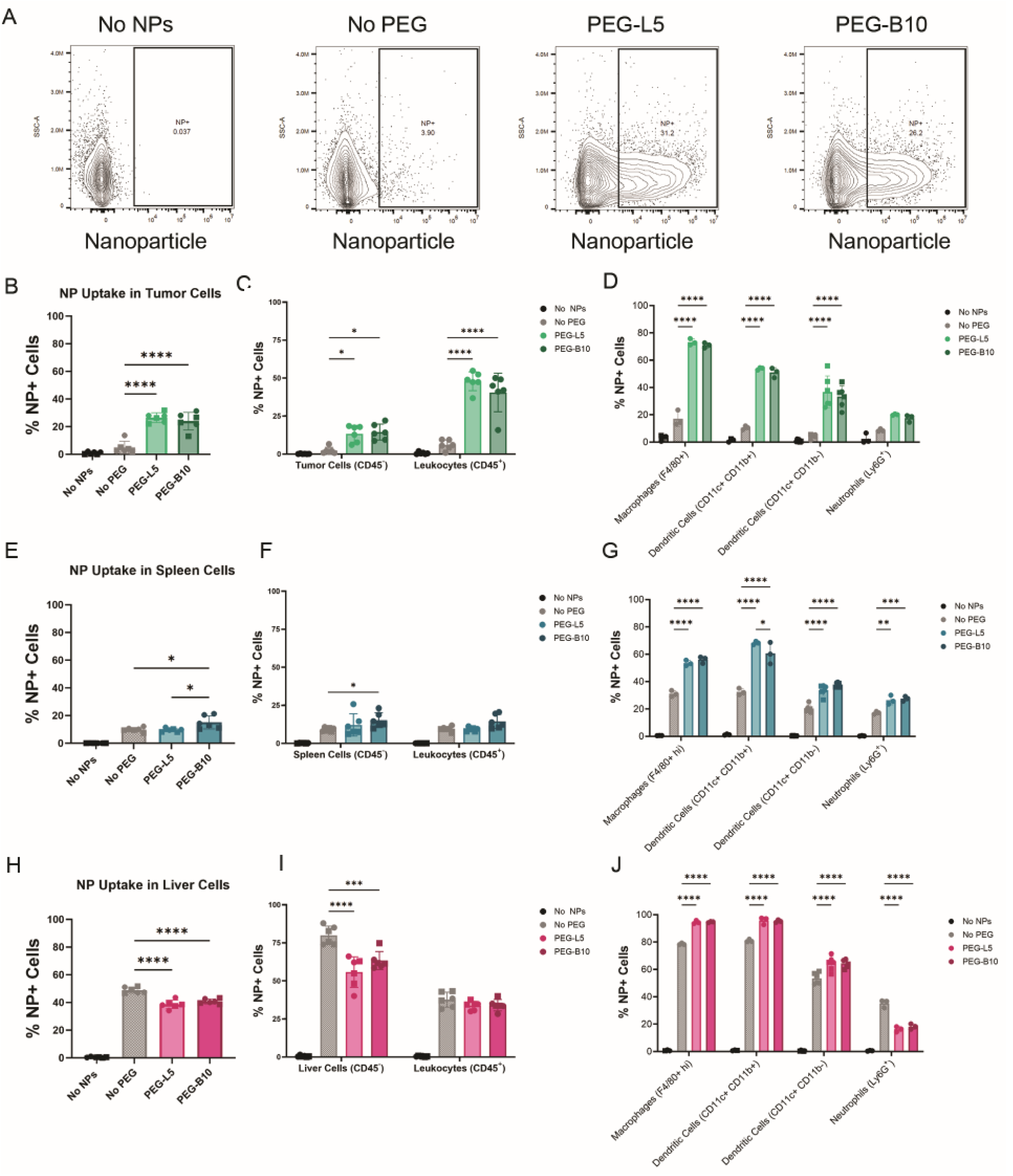
NP immune cell uptake in 4T1 tumor bearing balb/c mice. **(A)** Representative flow cytometry contour plots of NP gating strategy for live cells isolated from 4T1 tumors. Percent of **(B)** live cells, **(C)** leukocytes or tumor cells, **(D)** neutrophils, dendritic cells, or macrophages containing NP obtained from tumor tissue. Percent of **(E)** live cells, **(F)** leukocytes or CD45-spleen cells, **(G)** neutrophils, dendritic cells, or macrophages containing NPs obtained from spleen tissue. Percent of **(H)** live cells, **(I)** leukocytes or liver cells, **(J)** neutrophils, dendritic cells, or macrophages containing NPs obtained from liver tissue. *p<0.05, **p<0.01, and ***p<0.001 by two-way ANOVA with Tukey correction. N=6 mice for No PEG, PEG-L5, and PEG-B10, N=4 mice for No NPs

As in other models, PEG-B10 increased total uptake in spleen cells relative to linear and non-PEGylated (**Figure 7E and Figure S10C,D**), though only CD45-splenocyte uptake was significantly increased (**Figure 7F**). Both linear and branched PEGylation drove an increase in macrophage, DC, and neutrophil uptake compared to non-PEGylated. Non-PEGylated NPs had the highest uptake within the liver of 4T1 bearing mice, including a substantial percentage of CD45-liver cells (**Figure 7H-I and Figure S10E,F**). Macrophages and dendritic cells within the liver of the balb/c mice were the dominant phagocytes internalizing NPs. Both linear and branched PEGylation increased uptake to over 90% of macrophages and CD11b+ DCs, and was similar between PEGylation types (94.5-95.8%) (**Figure 7J**). However, both forms of PEGylation shielded NPs from neutrophil internalization (16.3-18.2%) reducing by roughly half of that seen for non-PEGylated (35.2%). The NP uptake patterns within the spleen were similar to the findings of the nude mice where the PEG-B10 NPs had the highest uptake, and the macrophages and dendritic cells had the highest uptake for all groups. However, DCs were differentially affected by PEGylation type in nude mice relative to wild type balb/c, which suggests that T lymphocytes play a role in NP uptake by phagocytes.

## Discussion

In this study, we examined how PEG architecture affected NP biodistribution in tumors originating from two different cell lines. We utilized 4T1 cells, a mouse mammary carcinoma, due to the extensive characterization of this cell in line in relation to NP infiltration of the tumor microenvironment. We also used A549 cells, which are a human lung adenocarcinoma cell line, to determine whether the type of cancer and its resulting tumor microenvironment affected the results. Further, 4T1 tumors were analyzed in both athymic nude mice and immune-competent wild type mice to determine the role of T cells, which are absent in nude mice similar to patients undergoing lymphodepletion during cancer therapy (32). Recent studies have indicated that branched PEG may offer enhanced NP circulation time as well as evasion of the accelerated blood clearance phenomenon (30). We previously found that PEG branching can increase NP mobility in the extracellular matrix (15) which could lead to greater NP accumulation within tumors where there is an upregulation of ECM components (35,36). Therefore, we utilized *in vivo* and *ex vivo* imaging to analyze the biodistribution of NPs coated with either linear 5 kDa PEG, 10 kDa 4-arm branched PEG, or no PEG and hypothesized that the branched PEG coated NPs would have a greater accumulation and retention at the tumor site compared to NPs coated with linear PEG or no PEG. Finally, we used flow cytometry to quantify cellular uptake by phagocytes within each tumor model to determine whether PEGylation strategy affected immune cell-specific internalization profiles.

Our imaging results indeed showed that branched PEG coated NPs had the highest accumulation in the 4T1 tumors compared to the other types of NPs. Furthermore, the non-PEGylated NPs had a distinctly high accumulation in the liver compared to other organs and other NP types which suggests rapid clearance from the body. However, in the A549 tumors, the linear PEG coated NPs had the highest NP accumulation although the differences were not statistically significant. The overall trends for where NPs accumulated depending on PEG coating were the same in both 4T1 and A549 mouse models where the non-PEGylated NPs mostly accumulated in the liver while PEGylated NPs had deposited mainly in the spleen followed by the liver and then tumor. Previous studies have shown that immune cells can play an important role in both transporting NPs to the tumor site leading to broader distribution within the tumors (21,23,37–39). Therefore, it is important to understand how NP coatings can affect both immune cell infiltration into tumors as well as uptake of NPs by different immune cells.

We found that the increased overall accumulation of both linear and branched PEGylated NP at the tumor site could be attributed in large part to their uptake by tumor associated phagocytes such as macrophages and dendritic cells. While PEGylation is often used to provide stealth function for cancer nanomedicines, it may also prove beneficial to identify PEGylation strategies which promote immune cell uptake within the TME. Interestingly, branched PEGylation appeared to affect immune cell uptake differently at the tumor site compared to other filtering organs, with the most notable differences observed in the spleen. Plausible explanations for leukocyte accumulation at these sites could be either (i) a unique protein corona signature for PEG-B10 NP or (ii) a PEG-dependent immune cell uptake mechanism. We have shown in prior work branched PEGylation can reduce protein adsorption (15), but have yet to characterize protein corona composition as a function of PEG structure. While we expect both PEG-L5 and PEG-B10 to possess dense PEG brushes on their surfaces (15), the branched conformation of PEG-B10 will likely alter interchain repulsion and entanglement within the polymer brush leading to altered NP surface morphology. For example, prior work has demonstrated how surface coatings assembled on NP using highly branched PEG led to a more rugged surface that better resists protein adhesion and enables greater uptake *in vitro* compared to traditional linear PEG (40). Together with our current findings, these results highlight the need for more comprehensive profiling of how PEG architecture influences immune cell uptake, both systemically and within tumors, given its potential broader implications for immune responses to PEGylated nanotherapeutics (41–43).

Neutrophil uptake in tumors and other tissues was an unexpected finding. The effect of NP PEGylation modulating uptake has been demonstrated in circulating neutrophils in other models (44,45), but the influence of neutrophilic NP uptake in a tumor setting is not as well understood. PEGylation increased neutrophil uptake within tumors to approximately 20% which may be therapeutically relevant. It should be noted that every tumor microenvironment is different as is the tumor’s effect on systemic hematopoeisis and generation of neutrophils. The 4T1 model has a well characterized increase in neutrophil presence that supports immunosuppression (sometimes referred to as granulocytic myeloid derived suppressor cells or low density neutrophils) (33). Indeed, we show high neutrophil presence in spleens in mice bearing 4T1 tumors as well as the tumor itself, and NP uptake within these cells. Such neutrophil uptake might be clinically relevant in tumor types with that are enriched for these tumor supporting neutrophil phenotypes.

Similarly, NP clearance by the MPS may not be an isolated system from the tumor microenvironment. The presence of a tumor can alter systemic leukocyte and stromal cell function in the liver and spleen, meaning nanoparticle clearance observed in healthy models may not reflect what occurs in a cancer model. Previous characterization of systemic immunomodulation by tumors have shown impairment of DC maturation, M1 macrophage polarization, and vaccine responses by T cells (46). This underscores the potential importance of studying the systemic clearance of PEGylated nanoparticles in tumor bearing models, where conventional clearance pathways may be altered and could present new therapeutic opportunities. For example, effective cancer vaccine loaded NPs are associated with systemic immunity and memory lymphocytes within the spleen. We found that primary BMDCs and the RAW macrophage cell line exhibited greater uptake of nanoparticles coated with branched PEG compared to those with linear PEG, consistent with the overall trend of increased splenic accumulation of PEG-B10 nanoparticles across the models tested in our study. Conversely, the *in vitro* uptake patterns of nanoparticles with either linear or branched PEG by phagocytes did not reflect the uptake behavior observed in tumor-associated phagocytes *in vivo*. However, this is not surprising and suggests that the effect of PEGylated NP uptake by phagocytes may be highly dependent on TME-driven factors rather than a universal response to PEGylation. It is also unknown whether these are the result of functional differences in tumor infiltrating cells or via indirect interactions with stromal cells in the TME. We also note other effects such as endothelial transport or altered diffusion through ECM may influence phagocyte uptake and future studies are required to determine these contributions.

## Conclusion

We found that branched PEGylation of NPs has a profound effect on biodistribution and uptake profile by phagocytes both within tumors and within filtering organs, such as the spleen, relative to linear PEGylation counterparts. PEGylation was necessary for accumulation within tumors, and NP uptake changes between branched and linear PEG were dependent on both tumor model and immune profile of the host animal. Overall, these findings offer valuable insight for optimization of NP delivery systems by tailoring PEGylation strategies to the specific target cell—such as dendritic cells, macrophages, or cancer cells—and desired site of action, whether within the TME or for systemic immune modulation.

## Acknowledgements

This Research was supported in part by: the Center for Cancer Research, National Cancer Institute, National Institutes of Health Intramural Research Program project number ZIA BC 012021; federal funds from the National Cancer Institute, National Institutes of Health, under contract HHSN261200800001E; the Cancer Innovation Laboratory; the NCI-UMD Partnership for Integrative Cancer Research; and the National Science Foundation (CAREER Award 2047794). The contributions of the NIH authors were made as part of their official duties as NIH federal employees, are in compliance with agency policy requirements, and are considered Works of the United States Government. However, the findings and conclusions presented in this paper are those of the author(s) and do not necessarily reflect the views of the NIH or the U.S. Department of Health and Human Services, nor does mention of trade names, commercial products, or organizations imply endorsement by the U.S. Government. S.M.K. and C.M.J. are employees of the Veterans Affairs Maryland Health Care System. The views reported in this paper do not reflect the views of the Department of Veterans Affairs or the United States Government. C.M.J. has equity positions with Cartesian Therapeutics, Barinthus Biotherapeutics, and Nodal Therapeutics.

