## Supplementary Information for "PEGylation strategies for enhanced nanoparticle delivery to tumor associated immune cells"

**Affiliations**: ^1^Fischell Department of Bioengineering, University of Maryland, College Park, MD 20742, USA; ^2^Cancer Biomaterials Engineering Section, Cancer Innovation Laboratory, Center for Cancer Research, National Cancer Institute, Frederick, MD, 21702 USA; ^3^Small Animal Imaging Program, Laboratory Animal Sciences Program, Leidos Biomedical Research, Inc., Frederick National Laboratory for Cancer Research, Frederick, MD 21701, USA; ^4^Fischell Institute of Biomedical Devices, University of Maryland, College Park, MD 20742, USA; ^5^Department of Veterans Affairs, Veterans Affairs Maryland Health Care System, Baltimore, MD 21201; ^6^Department of Microbiology and Immunology, University of Maryland School of Medicine, Baltimore, MD 21201; Marlene and Stewart Greenebaum Comprehensive Cancer Center, University of Maryland, Baltimore, MD 21201

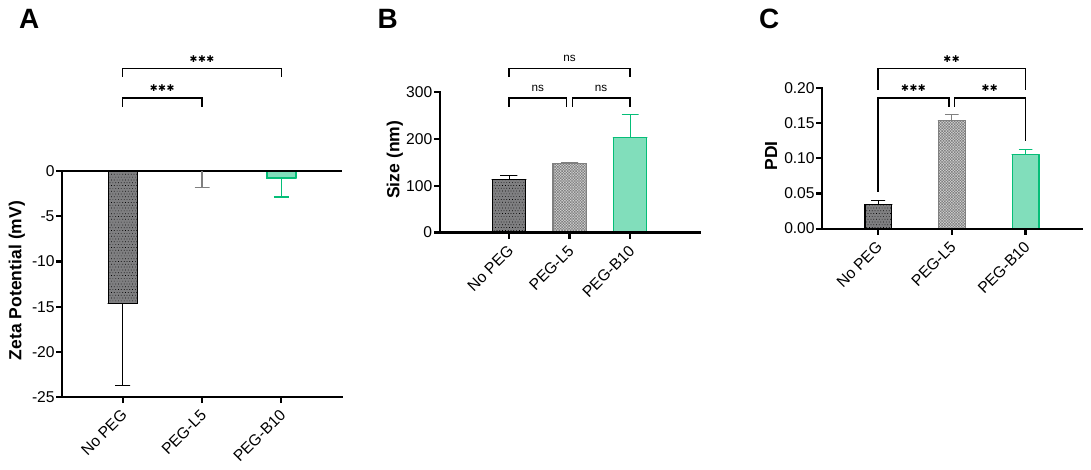

**Figure S1. Nanoparticle Characterization.** (A) Zeta potential of NIR NPs. (B) Hydrodynamic diameter of NIR NPs. (C) Polydispersity index (PDI) of NIR NPs. *p<0.05, **p<0.01, and ***p<0.001 by one-way ANOVA with Tukey correction; N=3.

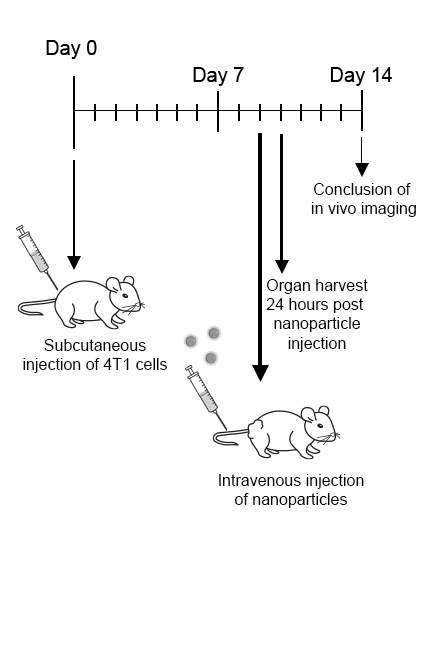

**Figure S2. 4T1 Injection Timeline.** Timeline showing the establishment of the 4T1 tumor model and subsequent NP injection and tissue harvest as well as the timeline for *in vivo* IVIS imaging.

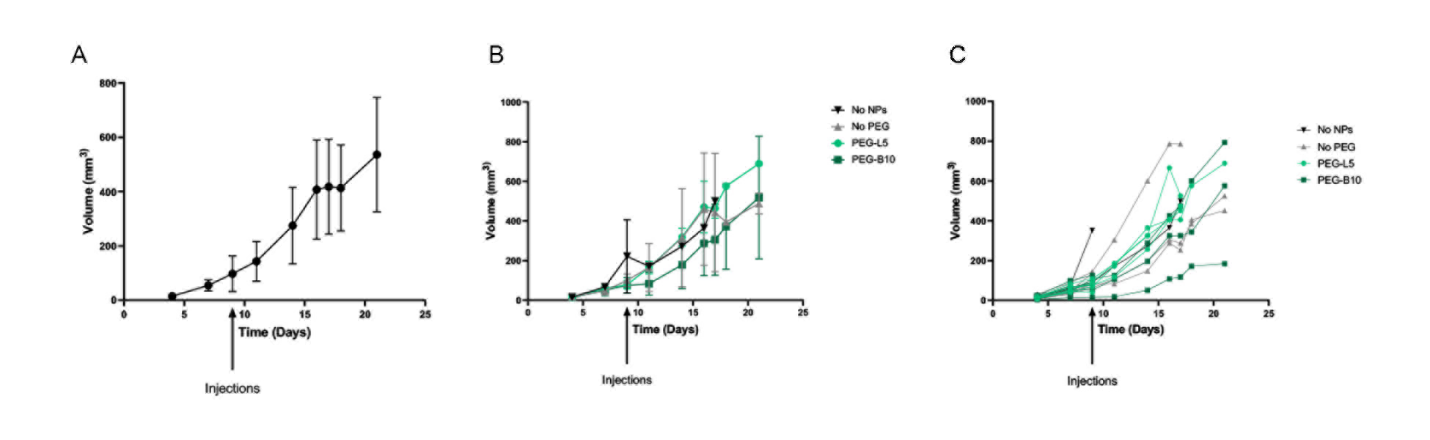

**Figure S3. 4T1 Tumor Growth in Nude Mice.** (A) Average tumor growth in 4T1 tumor bearing nude mice. (B) Average tumor growth in 4T1 tumor bearing nude mice according to treatment group. (C) Tumor growth for individual mice according for all treatment groups.

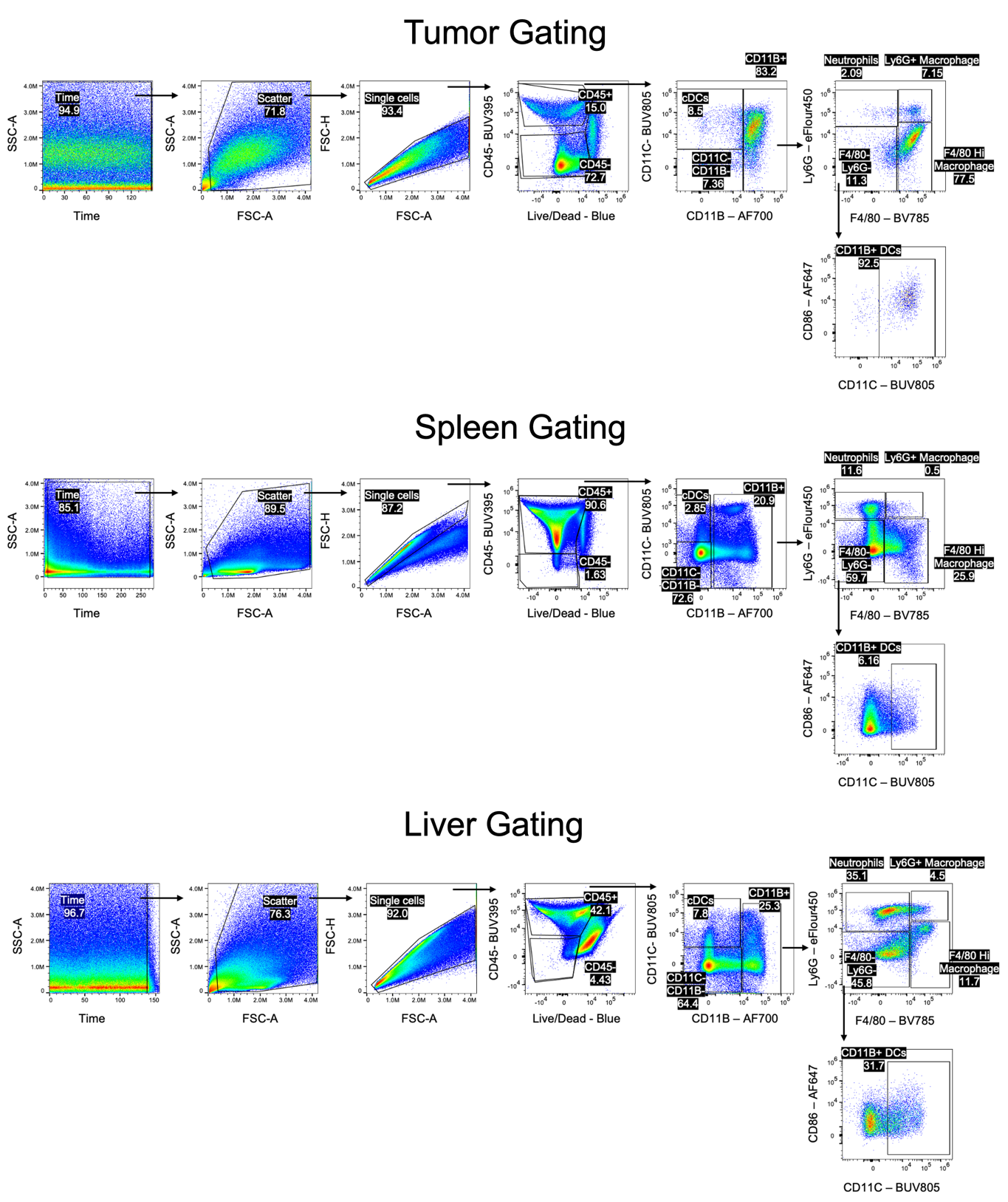

**Figure S4. 4T1 Gating strategy for the tumor, spleen and liver.** Representative flow cytometry plots showing the gating strategy used for identifying different immune cell populations in tumor, spleen and liver.

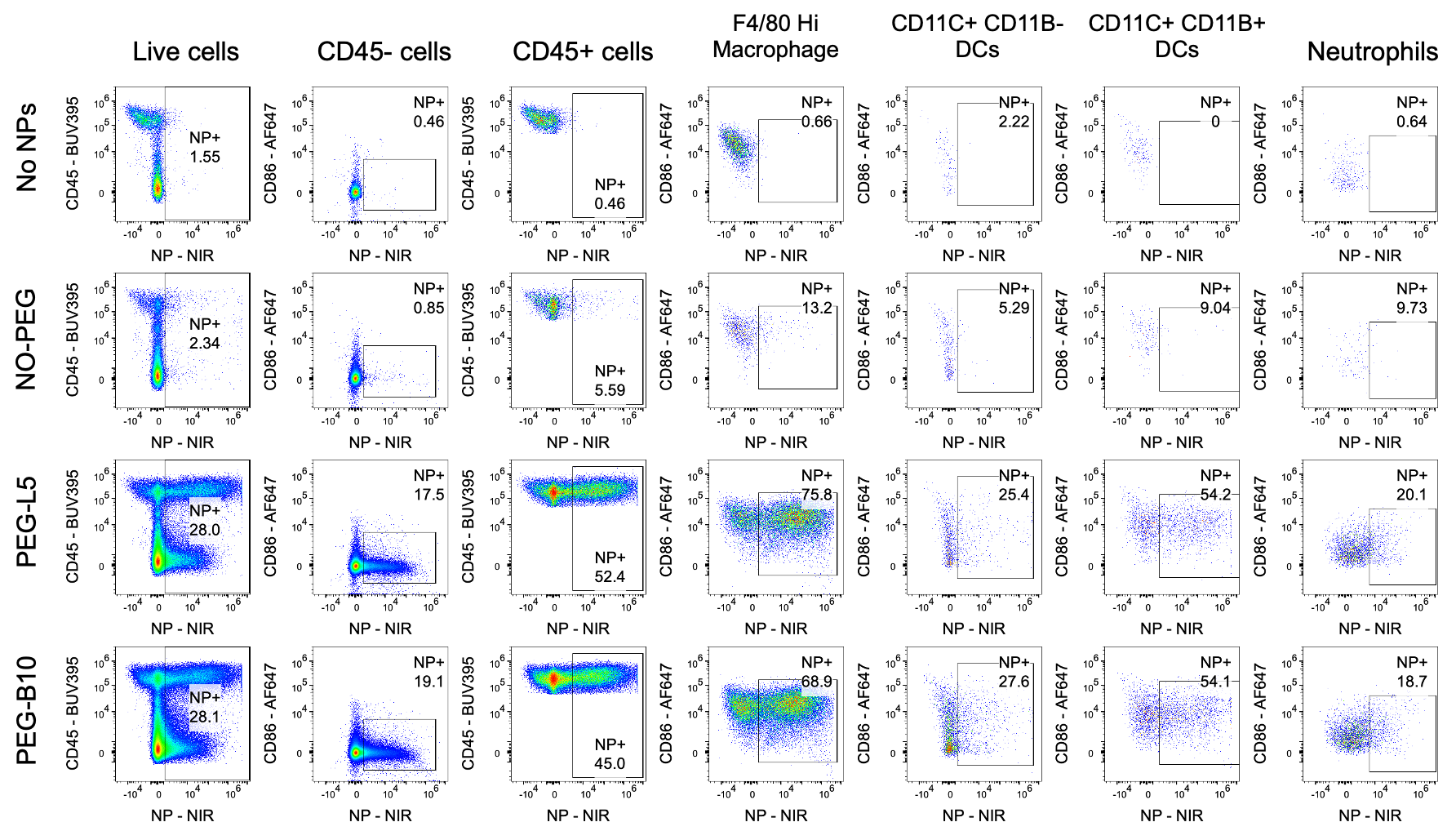

**Figure S5. Flow plots showing the NP+ cells:** Representative flow plots showing the NP+ cells from 4T1 Tumor bearing NUDE mice in different treatment groups.

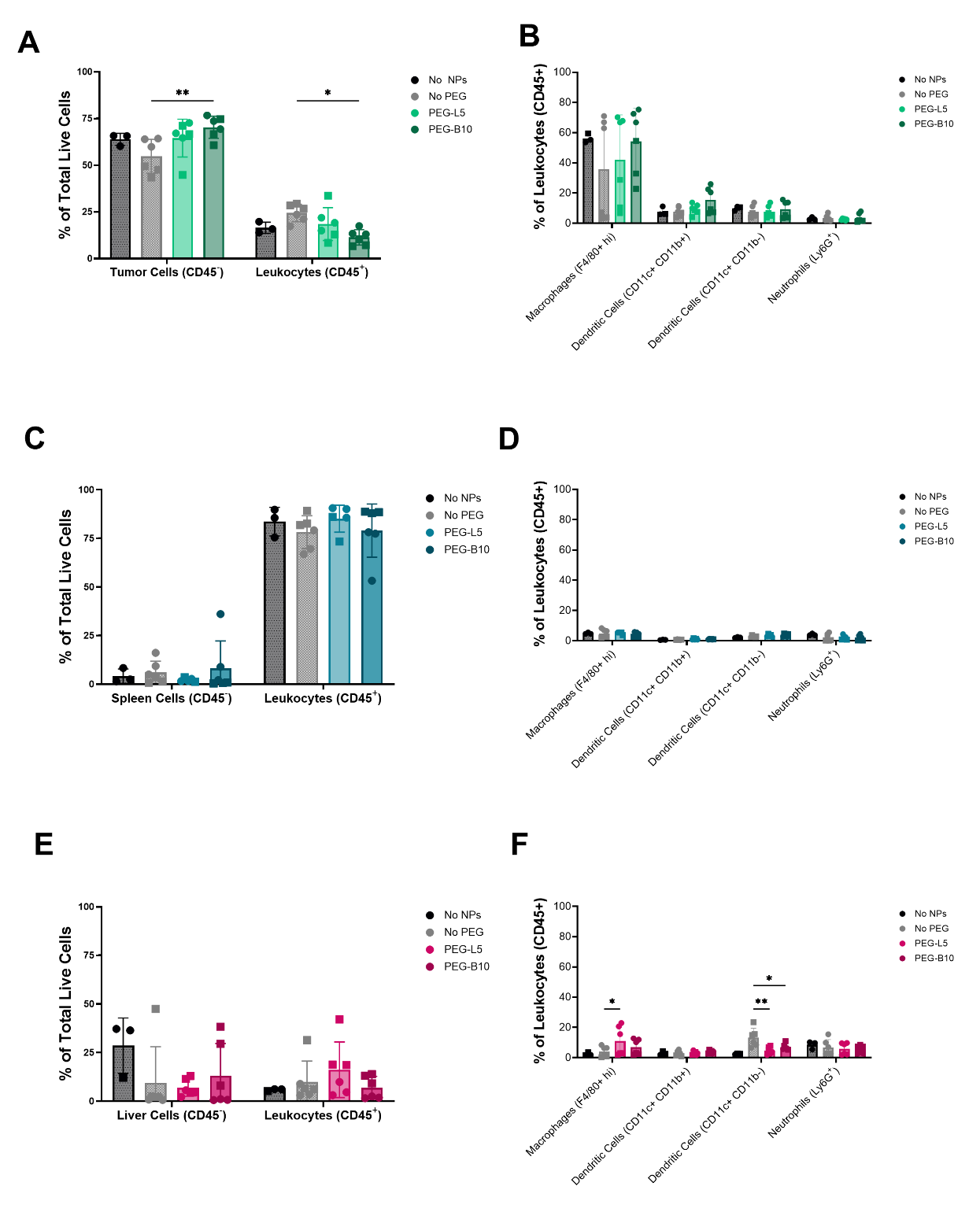

**Figure S6. 4T1 Tumor Bearing Nude Mice Immune Cell Composition.** (A) Distribution of CD45- tumor cells and CD45+ leukocytes in 4T1 tumors. (B) Distribution of macrophages, CD11b+ DCs, CD11b- DCs, and neutrophils in 4T1 tumors. (C) Distribution of CD45- spleen cells and CD45+ leukocytes in spleen tissue collected from 4T1 tumor bearing nude mice. (D) Macrophage, CD11b+ DC, CD11b- DC, and neutrophil distribution in spleen tissue collected from 4T1 tumor bearing nude mice. (E) CD45- liver cell and CD45+ leukocyte distribution in liver tissue collected from 4T1 tumor bearing nude mice. (F) Distribution of macrophages, CD11b+ DCs, CD11b- DCs, and neutrophils in liver tissue collected from 4T1 tumor bearing nude mice. *p<0.05, **p<0.01, and ***p<0.001 by two-way ANOVA with Tukey correction. N=6 mice for No PEG, PEG-L5, and PEG-B10, N=4 mice for No NPs

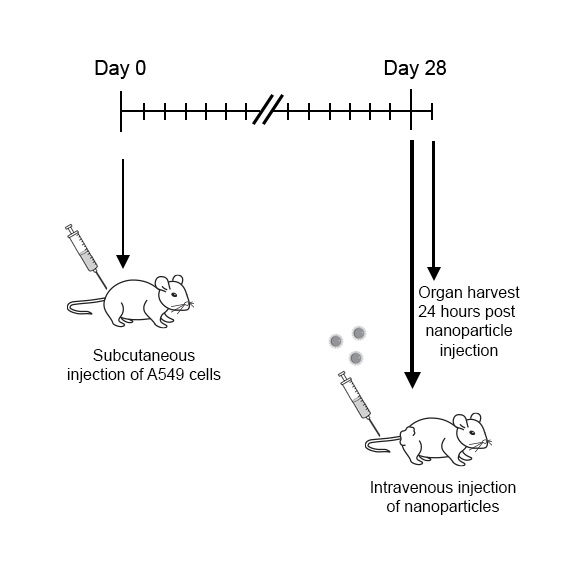

**Figure S7. A549 Injection Timeline and Tumor Growth.** Timeline showing the establishment of the A549 tumor model and subsequent NP injection and tissue harvest.

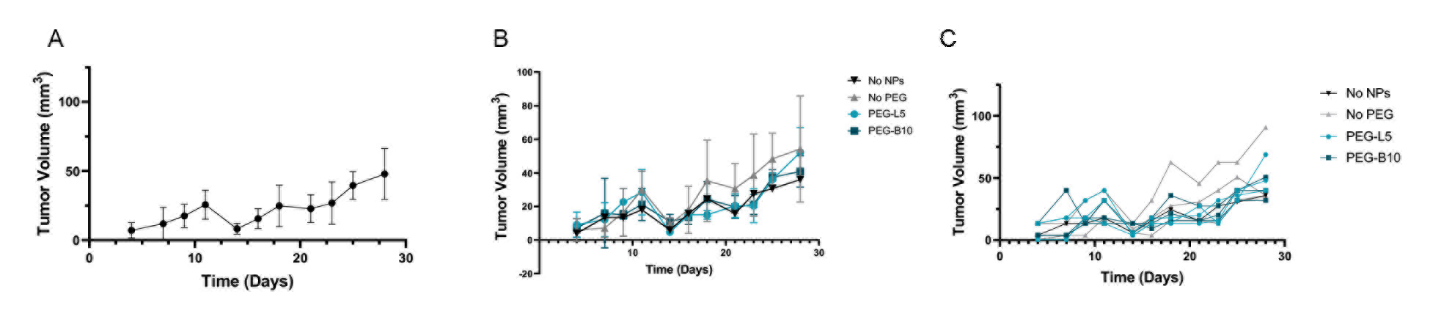

**Figure S8. A549 Injection Timeline and Tumor Growth in Nude Mice.** (A) Average tumor growth in 4T1 tumor bearing nude mice. (B) Average tumor growth in 4T1 tumor bearing nude mice according to treatment group. (C) Tumor growth for individual mice according for all treatment groups.

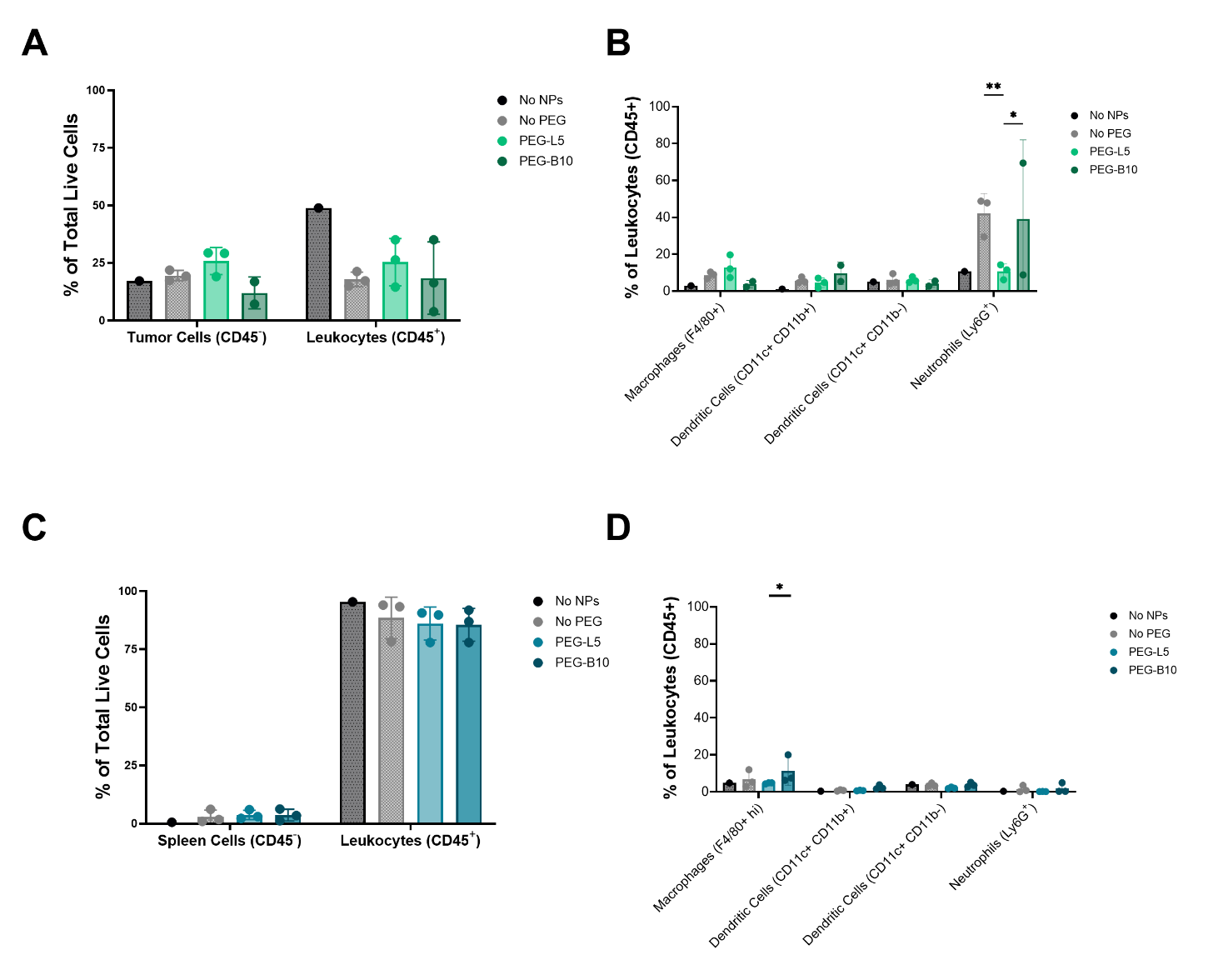

**Figure S9. A549 Tumor Bearing Nude Mice Immune Cell Composition. (**A) Distribution of CD45- tumor cells and CD45+ leukocytes in A549 tumors. (B) Distribution of macrophages, CD11b+ DCs, CD11b- DCs, and neutrophils in A549 tumors. (C) Distribution of CD45- spleen cells and CD45+ leukocytes in spleen tissue collected from A549 tumor bearing nude mice. (D) Macrophage, CD11b+ DC, CD11b- DC, and neutrophil distribution in spleen tissue collected from A549 tumor bearing nude mice. *p<0.05, **p<0.01, and ***p<0.001 by two-way ANOVA with Tukey correction. N=3 mice for No PEG, PEG-L5, and PEG-B10, N=1 mouse for No NPs.

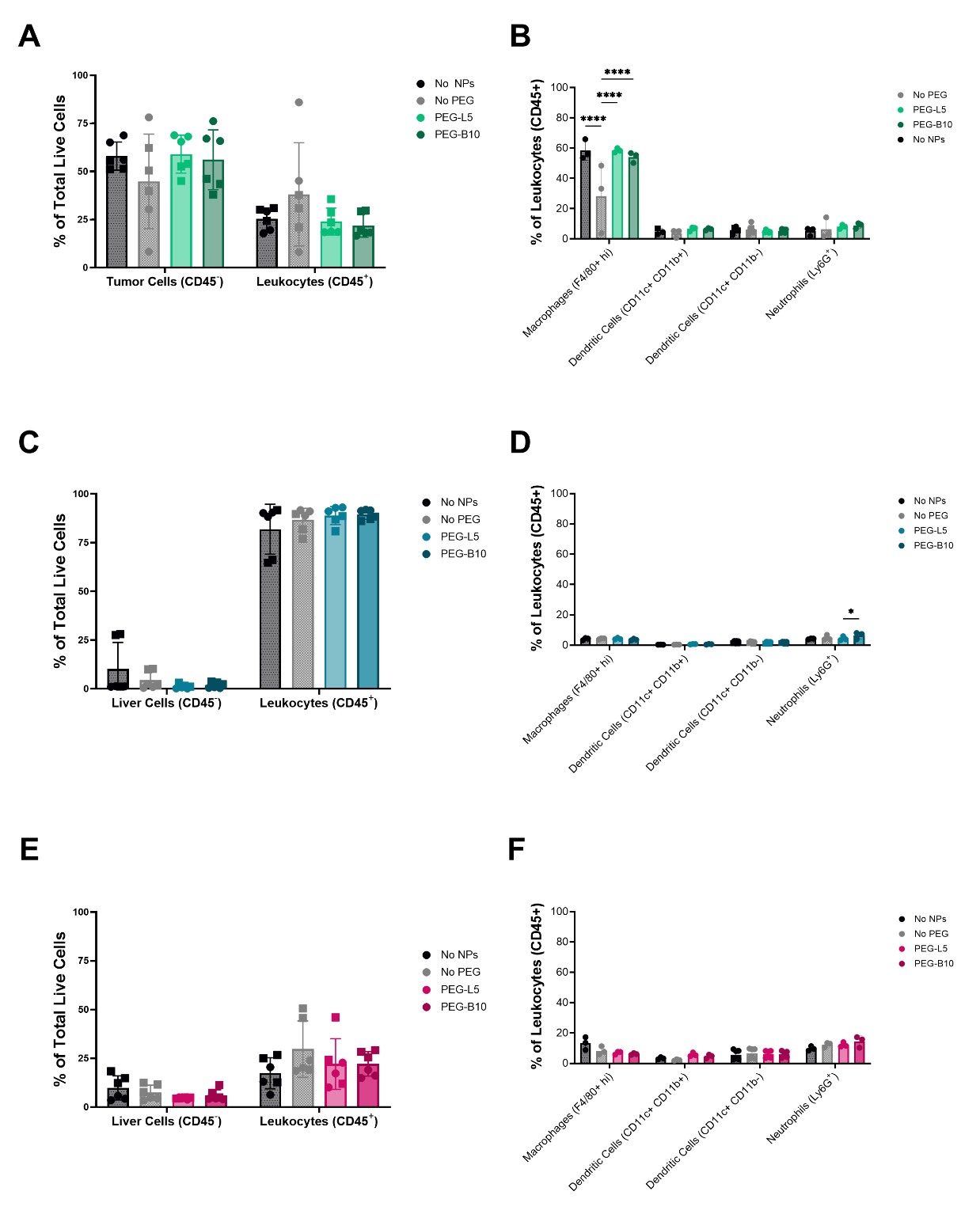

**Figure S10. 4T1 Tumor Bearing Balb/c Mice Immune Cell Composition.** (A) Distribution of CD45- tumor cells and CD45+ leukocytes in 4T1 tumors in balb/c mice. (B) Distribution of macrophages, CD11b+ DCs, CD11b- DCs, and neutrophils in 4T1 tumors in balb/c mice. (C) Distribution of CD45- spleen cells and CD45+ leukocytes in spleen tissue collected from 4T1 tumor bearing balb/c WT mice. (D) Macrophage, CD11b+ DC, CD11b- DC, and neutrophil distribution in spleen tissue collected from 4T1 tumor bearing balb/c WT mice. (E) CD45- liver cell and CD45+ leukocyte distribution in liver tissue collected from 4T1 tumor bearing balb/c WT mice. (F) Distribution of macrophages, CD11b+ DCs, CD11b- DCs, and neutrophils in liver tissue collected from 4T1 tumor bearing balb/c mice.
